# Clinical characteristics, drug-resistance genes and virulence analysis of carbapenem-resistant Klebsiella pneumoniae

**DOI:** 10.1101/2024.10.25.620218

**Authors:** Junning He, Peilin Zhang, Cheng Li, Ning Xie, Yijie Yin, Guixia Wang, Xianting Liang, Yongfang Liu

## Abstract

**OBJECTIVE:** The aim was to investigate the clinical infection characteristics, risk factors and convergence of carbapenem-resistant and hypervirulent Klebsiella pneumoniae(hv-CRKP) in our hospital, in order to provide scientific basis for the rational use of antibiotics in the clinic.

**METHODS:** CRKP strains were isolated from clinical specimens of 44 patients in central intensive care unit (CICU), neurosurgery intensive care unit (NICU) and emergency intensive care unit (EICU) of our hospital in 41 months. The clinical manifestations of the patients were analyzed, and the hypermucoviscous phenotype was defined by string test. The drug resistance gene, virulence gene and capsular specific serotype of the isolated strains were detected by polymerase chain reaction (PCR) technology, and the gene of CRKP producing KPC enzyme was sequence.

**RESULTS:** The incidence of sepsis in 44 patients with CRKP infection was 25.0%, and the overall treatment failure rate was 38.6%. All the 44 strains of CRKP produced KPC enzyme, and all of them were bla_KPC-2_ (100%);14 strains (31.8%) were positive in string test; 23 strains (52.3%) were found to have virulence related genes, including 10 strains (22.7%) of rmpA, 16 strains (36.4%) of iucA and 19 strains (43.2%) of peg-344. 5 strains (11.4%) were capsule-specific serotypes and 13 strains (29.5%) were identified as hv-CRKP.

**CONCLUSION:** The mechanism of CRKP resistance in our hospital is *bla*_KPC-2_, the detection rates of virulence gene, string test positive and hypervirulent capsular specific serotype are all relatively high in CRKP, there are not a few Klebsiella pneumoniae with carbapenem-resistant and hypervirulent.

## Introduction

*Klebsiella pneumoniae* (KP) is a common pathogen of respiratory infection as well as a serious pathogen of community-acquired infections. Carbapenems are β-lactam antibacterials with the broadest antimicrobial spectrum and the strongest antimicrobial activity to date, and they are one of the commonly used drugs for the treatment of Klebsiella pneumoniae infection. Antibacterials abuse and plasmid-mediated transfer of drug-resistant genes have led to the development of drug resistance in *Klebsiella pneumoniae*, and the emergence of carbapenem-resistant *Klebsiella pneumoniae* (CRKP) has led to a reduction in clinical efficacy. Genetic variation, expression of virulence genes, and changes in regulatory mechanisms lead to the production of hypervirulent *Klebsiella pneumoniae* (hvKP), these strains are more pathogenic and aggressive than classical *Klebsiella pneumoniae* (cKP)^[1]^, and are more likely to cause severe invasive and disseminated infections^[2,3]^. Carbapenem resistance and hypervirulence have become two different evolutionary directions of *Klebsiella pneumoniae*^[4]^ posing a great threat in the clinic. Carbapenem resistance and hypervirulence have long been recognized as two non-overlapping branches in different genetic contexts. However, in recent years, an increasing number of Klebsiella pneumoniae have been identified as integrating both phenotypes^[4]^, and this dual challenge makes the carbapenem-resistant and hypervirulent *Klebsiella pneumoniae* (hv-CRKP) has become the focus of nosocomial infection prevention and control, causing serious medical challenges and public health problems. The aim of our research was to study and analyze the resistance and virulence mechanisms of the clinically isolated CRKP strains in our hospital, define the high viscosity phenotype by string test^[5]^, detect resistance gene, virulence gene, and capsular serotype by PCR, and perform gene sequencing on all the KPCase-producing CRKP, so as to provide a scientific basis for the prevention and control of infection and drug decision-making of hv-CRKP.

### 1. Materials and Methods

### 1.1 Strain source

44 strains of CRKP were isolated from the Center ICU (CICU), Neurosurgery ICU (NICU) and Emergency ICU (EICU) of the Affiliated Hospital of North Sichuan Medical College from June 1, 2020 to October 31, 2023 were enrolled in the study. Inclusion criteria: (1) CRKP infection occurred in the lung or other parts of the body during ICU admission, and infection was defined as the invasion of CRKP into the patient’s tissues of various systems and the corresponding clinical manifestations; (2) The strain was resistant to any of the carbapenems of imipenem, meropenem, ertapenem; (3) To prevent data duplication, only the first positive specimen of infection in the same patient was included. A total of 44 clinically isolated strains of CRKP were isolated from sputum, blood or urine specimens of 44 patients. The 44 patients and 44 strains of CRKP were numbered as cases 1 to 44 and strains 1 to 44, respectively. The study has been approved by the Ethics Committee.

### 1.2 Methods

#### 1.2.1 Resistance gene detection

The 44 strains of CRKP from clinical specimens were tested for resistance genes, and the primers were synthesized by Shanghai Sangon Biotech, as shown in Table 1. The strains kept in the refrigerator at -80°C were recovered and inoculated on blood plates, incubated at 37°C for 24 hours, and 2-3 single colonies were selected with an inoculating loop into EP tubes containing 300 μl sterile distilled water. The bacterial suspension was prepared by mixing thoroughly with a vortex shaker, and genomic DNA was extracted by boiling method, the DNA template was obtained from the supernatant after centrifugation at 12,000 rpm for 10 min. PCR method was used to detect five carbapenemase-resistant genes, which include *bla*_KPC_, *bla*_NDM_, *bla*_IMP_, *bla*_VIM_ and *bla*_OXA-48_. Positive strains of *Escherichia coli* ATCC25922 and *Klebsiella pneumoniae* ATCC 700603 (National Center for Medical Culture Collections) are known to carry the target genes, and negative strains (carbapenem-sensitive strains detected from clinical specimens) without the target genes were used as controls.

**Table 1.**
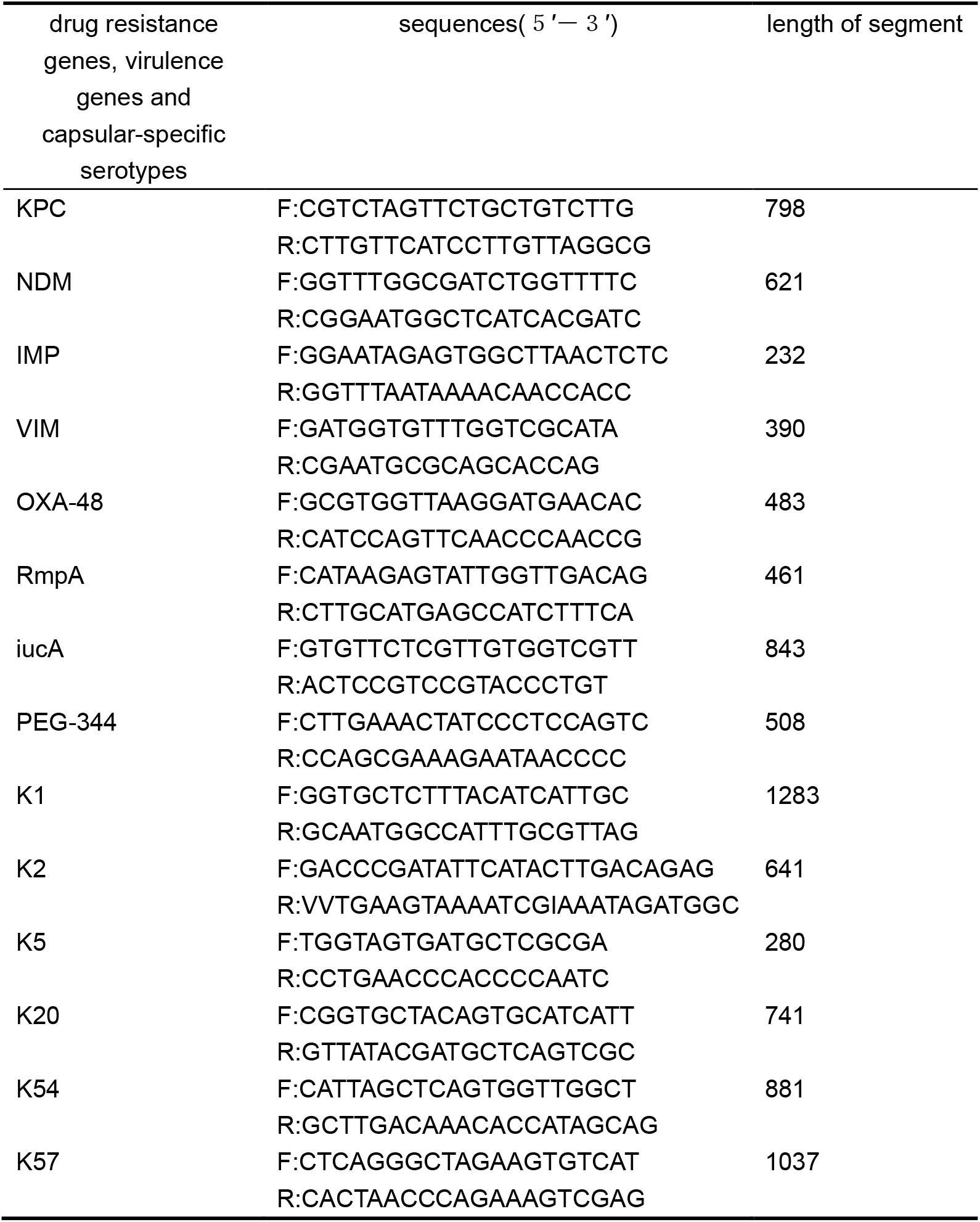
Sequence of common drug resistance genes, virulence genes and capsular-specific serotypes.

#### 1.2.2 Sequencing of KPC subtype

Genomic DNA extracted from 44 CRKP strains, and primers were designed to cover the full length of the *bla*_KPC_ gene according to the Genbank (www.ncbi.nlm.nih.gov/blast/). The extracted genomic DNA was amplified by PCR, and the purified PCR products were sequenced by sanger sequencing. The sequencing results were mapping with the *bla*_KPC-2_ sequences reported in Genbank, and mutations and subtype-specific sites were identified.

#### 1.2.3 The string test

44 CRKP strains were extracted from the frozen state and standardized inoculated on blood plates, incubated at 37°C for 24 hours, and suitable colonies were selected for string test. The bacterial colonies were stretched on the blood plate by stretching ring to observe whether they formed mucoid filament with a length of >5 mm,and if the filament was >5 mm, it was judged that the string test was positive, that is, hypermucoviscous phenotype.

#### 1.2.4 Detection of virulence genes and capsular specific serotype

Genomic DNA was extracted from 44 CRKP strains and three common virulence genes, *rmpA,iucA* and *peg-344*, as well as six common capsular serotype specific genes *K1, K2, K5, K20, K54* and *K57* were amplified by PCR. The primers were synthesized by Shanghai Sangon Biotech, as shown in Table 1 for details.

#### 1.2.5 Clinical data collection

Clinical information of the 44 patients with CRKP infection were collected, including demographic characteristics (name, gender, age), departmental distribution, admission and exit time, underlying disease (tumor, hypertension, diabetes mellitus, renal insufficiency, immunodeficiency, etc.), invasive operations (tracheal intubation, surgery, etc.), source of specimens, prognosis, etc.

## 2. Results

### 2.1 Clinical features of patients

44 patients were included in this study, 33 males and 11 females, aged 17-82 years, among whom 38 (86.4%) had nosocomial infection and 6 (13.6%) had community infection. The main source of patients was NICU (52.3%, 23/44), followed by CICU (38.6%, 17/44) and EICU (9.1%, 4/44). All 44 patients (100%) underwent invasive operations such as endotracheal intubation and manual surgery. Sepsis occurred in 11 patients (25.0%) and septic shock occurred in 9 patients (20.5%). The patients with underlying diseases before admission were hypertension (50.0%, 22/44), type 2 diabetes mellitus (29.5%, 13/44) and tumor (2.3%, 1/44). The prognosis of patients was as follows: 27 cases (61.4%) were discharged after infection improved, 14 cases (31.8%) were discharged automatically after infection did not improve, 3 cases (6.8%) died, and the treatment failure rate was 38.6%. The main specimens were sputum (79.5%, 35/44), and other specimens were from blood (9.1%, 4/44), urine (4.5%, 2/44) ascites (2.3%, 1/44), wound secretions (2.3%, 1/44) and cerebrospinal fluid (2.3%, 1/44).

### 2.2. Detection of drug resistance genes

*bla*_KPC_ was detected in all 44 CRKP strains (100%), and *bla*_NDM_,*bla*_IMP_,*bla*_VIM_ and *bla*_OXA-48_ were not detected (Fig 1).

**Figure 1.**
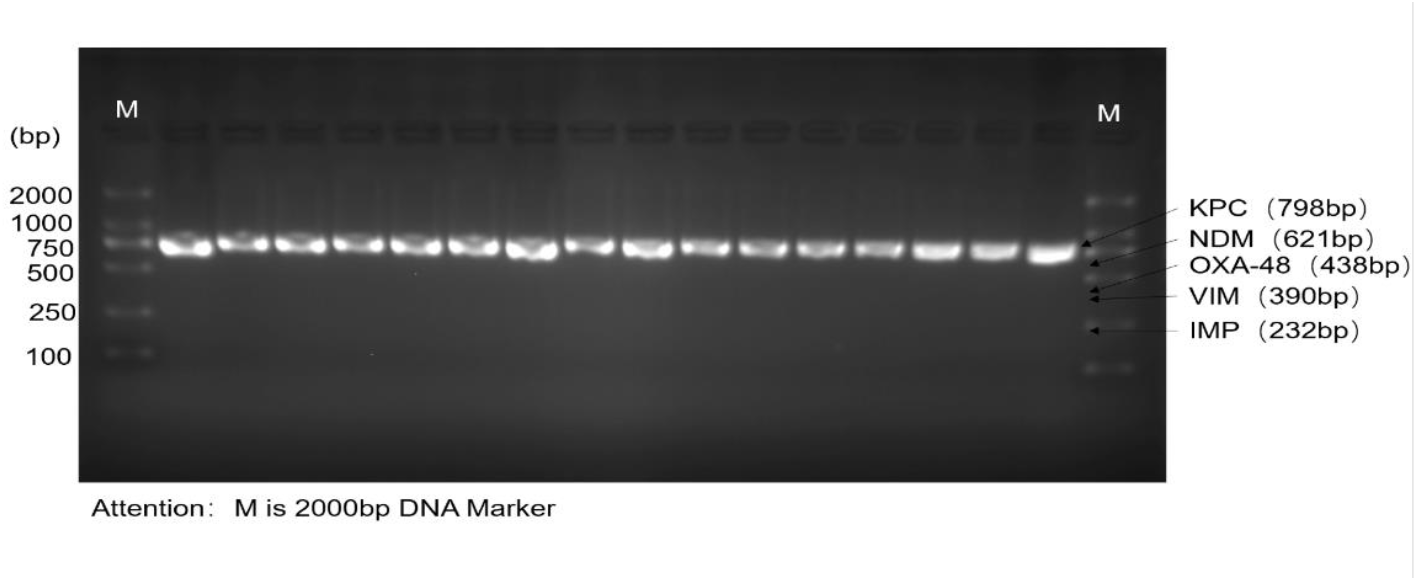
Electrophoretic PCR amplification of some KPC resistance genes

### 2.3 Sequencing of KPC subtypes

All PCR positive products were sequenced and analyzed by Sanger sequencing. 44 CRKP strains carrying *bla*_KPC_, all of which were KPC-2 (100%) subtype, and no other KPC subtypes were detected.

### 2.4 The string test

At 37°C, 14 (31.8%) out of 44 CRKP isolates showed significant positive string test. Although the length of filamentation varied between different strains, it was in the range of 1-3 cm.

### 2.5 Detection of virulence genes

A total of 23 strains (52.3%) of the 44 CRKP strains amplified virulence genes, including 10 strains (22.7%) of *rmpA*, 16 strains (36.4%) of *iucA*, 19 strains (43.2%) of *peg-344* and four strains (9.1%) of any two virulence genes, 9 strains (20.5%) of three virulence genes. A total of 13 strains (29.5%) contained two or more virulence genes (Fig 2).

**Figure 2.**
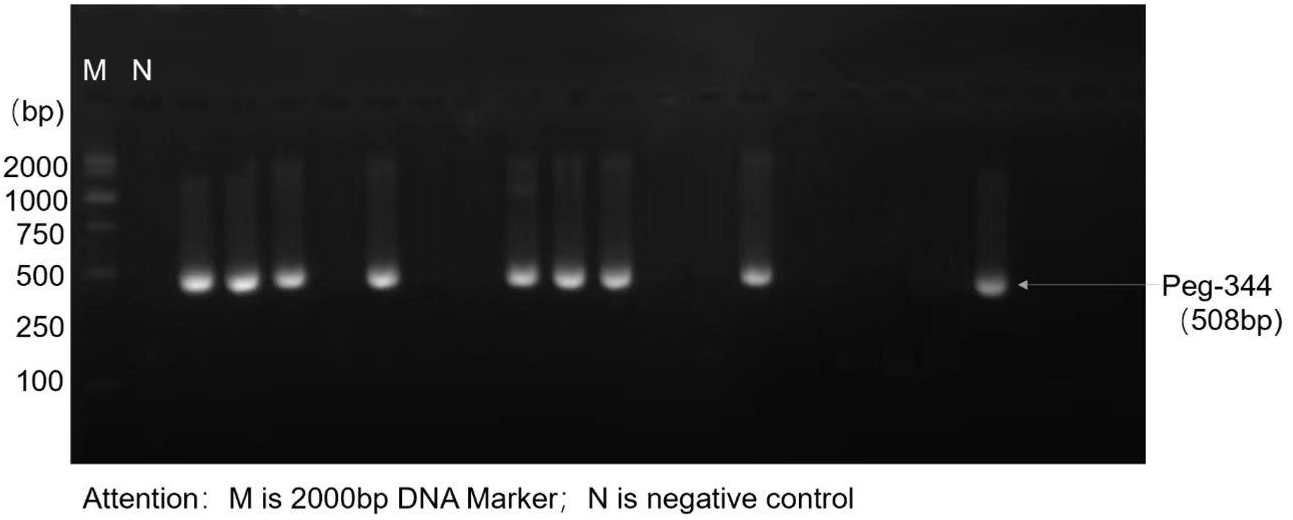
Electrophoretic PCR amplification of some peg-344 drug resistance gene

### 2.6 Detection of capsular specific serotype

Of the 44 CRKP strains, five (11.4%) amplified capsular specific serotypes, including one (2.3%) of type K1 was derived from the blood specimen of case 8, two (4.5%) of type K2 were derived from sputum specimens of cases 40 and 41, respectively, one (2.3%) of type K5 and one (2.3%) K57 from sputum specimens of cases 38 and 32, respectively. All five isolates were hv-CRKP and types K20 and K54 were not detected (Fig 3).

**Figure 3.**
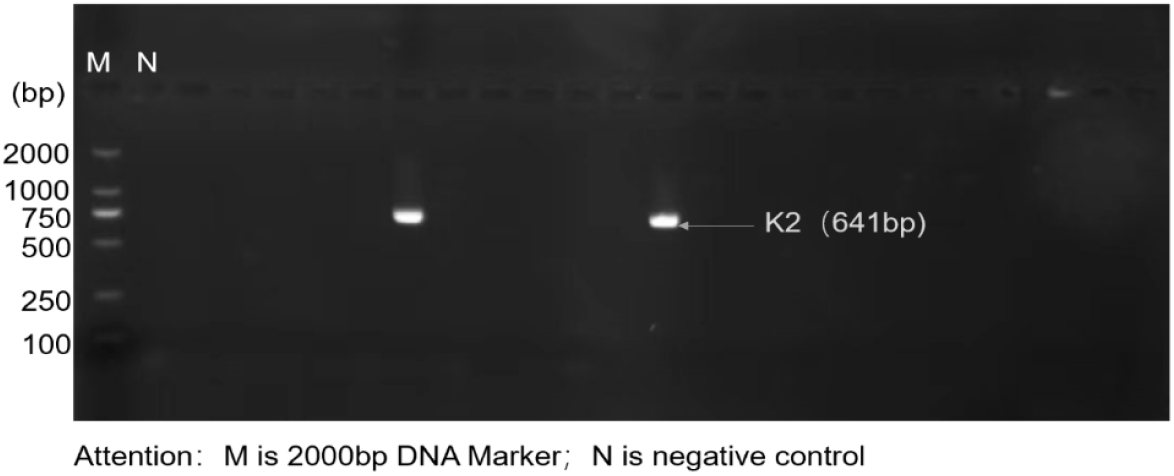
Electrophoretic PCR amplification of some K2 capsulor-specific serotype

## 3. Discussion

*Klebsiella pneumoniae* is an opportunistic pathogenic pathogen. Originally, carbapenem resistance and hypervirulence were considered to be two branches in different genetic backgrounds. However, with the horizontal transmission of plasmids carrying carbapenem resistance genes or virulence-encoding genes among different strains, carbapenem resistance and hypervirulence have gradually converged, and the difficulty of clinical treatment and infection prevention and control of *K. pneumoniae* has greatly increased. By summarizing and analyzing the current studies, the generation of hv-CRKP may occur through the following three pathways^[6]^: (1) CRKP acquired a hypervirulent phenotype; (2) hvKP acquired a carbapenem-resistant phenotype; and (3) cKP acquired a hybrid plasmid for carbapenem resistance and hypervirulence.

The drug resistance mechanisms of CRKP are complex and diverse, among which the most important is the production of carbapenemases. Studies have shown that *Klebsiella pneumoniae* carbapenemase (KPC) is the most common class A carbapenemase of CRKP in China^[7,8]^, whereas *NDM, IMP, VIM*, and *OXA-48* are more prevalent in some area of foreign countries. A KPC-2-producing *Klebsiella pneumoniae* strain was detected for the first time in a university hospital in Zhejiang, China in 2007^[9]^, and since then, the KPC-2-producing Enterobacteriaceae isolates began to spread rapidly and widely from that region. The current domestic and international studies on drug-resistant genotype KPC carried by CRKP strains showed that the predominant genotype in the United States was *bla*_KPC-3_ ^[10]^, while blaKPC-2 was the most common genotype in China^[11]^. In this experiment, all 44 CRKP strains carried KPC genes, and none of them carried *bla*_NDM_, *bla*_IMP_, *bla*_VIM_, *bla*_OXA-48_ genes. Sequencing results showed that all strains were *bla*_KPC-2_ (100%), which was basically consistent with the results of the current domestic and international studies,and no other mutant KPC subtypes were detected in this experiment. This study found that the detection rate of KPC carbapenemase in all strains belonging to both CRKP and hvKP was 100% and all of them were *bla*_KPC-2_, which was also detected 100% *bla*_KPC-2_ in our research, suggesting that the plasmid carrying blaKPC-2 may play an important role in the evolution of hv-CRKP.

HvKP often leads to highly pathogenic and severe infections in patients with autoimmune and even healthy populations, the most common being liver abscess, and also frequent multisite metastatic infections^[12,13]^, which are the main clinical features of hvKP. Current studies focus more on the virulence mechanism of hvKP. Although there is no precise and uniform criterion for the definition of hvKP, and there is still a lack of highly specific and sensitive markers, a variety of phenotypes and biomarkers have been characterized as criteria for the definition of hvKP^[14,15]^, such as hyperviscosity phenotypes, capsular specific serotype, virulence genes, and aerobic actin. Some scholars ^[16,17]^ believe that hvKP can be identified by genotype characteritics, with one or more virulence genes as the criteria for identification of hvKP, while others believe^**Error! Reference source not found**.[5]^ that hvKP can be identified by fulfilling at least two of the following criteria: (1) a positive string test; (2) a positive rmpA; and (3) a positive aerobic actin. Aerobic actin is the main siderophores produced by hvKP, and the siderophore aerobic actin has been identified as a factor that increases the virulence of hvKP compared with cKP^[5,18]^. Other studies have shown that pLVPKlike virulence plasmid plays an important role in the high viscosity expression and in the production of iron carriers in hvKP isolateds, which contains virulence-related determinants that encode regulators of mucus expression (rmpADC and rmpA2), iron carrier production (iutA iucABCD encoding aerobactin and the iroBCDN cluster encoding salmonellin), and metabolite transporter protein peg-344. Based on the definition criteria of most scholars, our research adopted simultaneous carrying of two or more virulence genes from *rmpA, iucA* and *peg-344* as the criterion for the standard for identifying hvKP. In our study, 10 strains (22.7%) of *rmpA*, 16 strains (36.4%) of *iucA*, and 19 strains (43.2%) of *peg-344* the 44 CRKP strains was identified both carbapenem resistance and hypervirulence. The high percentage of hv-CRKP suggests the importance of epidemiologic surveillance of this strain and enhanced infection prevention and control.

*Klebsiella pneumoniae* often exhibits a hyperviscous phenotype due to the production of a large amount of capsular polysaccharide (CPS) or the presence of specific virulence genes such as *rmpA*, which is defined by the positive string test. In this study, a total of 14 (31.8%) strains were positive in string test. For a long time, hyperviscosity has been regarded as equivalent to hypervirulence; however, new evidence has shown that not all hvKP strains are hyperviscous^[19]^, and some non-virulent *Klebsiella pneumoniae* also have a hyperviscous phenotype. In this study, one CRKP strain that did not carry any of the virulence genes of *rmpA, iucA, peg-344* and capsular specific serotypes was still positive in string test, which was not sufficient to determine whether the strain was hypervirulent or not. Therefore, phenotypes such as virulence genes or capsular serotypes need to be considered in the identification of hvKP, and positive string test can be used as a method to assist in the identification of hypervirulence.

hvKP has diverse capsular specific serotypes, up to now, 82 serotypes have been identified, the dominated ones being K1 and K2^[20]^, and other serotypes such as K5, K20, K54 and K57 were often reported to be associated with the virulence of *Klebsiella pneumoniae*. In this study, K1, K2, K5 and K57 were detected in 1 strain (2.3%), 2 strain (4.5%), 1 strain (2.3%) and 1 strain (2.3%), respectively. These 5 isolates were all hv-CRKP, and case 8 carrying type K1, cases 40 and 41 carrying type K2, developed severe pulmonary infection and septicemia 48 hours after admission. A study from Singapore and Taiwan^[21]^ demonstrated that capsular serotypes K1 or K2 had a more important impact in determining the virulence of *Klebsiella pneumoniae* than rmpA-positive non-K1/K2 strains, often showing greater virulence and pathogenicity, and were associated with severe pneumonia and sepsis.

In this study, the sources of strains were mainly NICU, CICU and EICU, and the common characteristics of hospitalized patients in such departments were that most of them were critically ill, with low immunity and have invasive operations. *Klebsiella pneumoniae* infection particularly affects people with weakened immune systems, such as the elderly and patients with chronic diseases or immunodeficiencies. Long-term use of Antibacterials, ventilator-assisted respiration and other invasive medical operations also increase the risk of infection with drug-resistant bacteria. In this study, 38 (86.4%) of 44 cases of CRKP were nosocomial infections with poor prognosis, sepsis rate up to 25%, and high therapeutic failure rate up to 38.6%, which should be highly concerned.

Currently, more and more hv-CRKP has been reported^[7]^. The resistance of hv-CRKP to conventional antimicrobials and carbapenem antimicrobials led to the difficulty of infection control, and the limitations of clinical selection of antimicrobials increased dramatically. Hypervirulence leads to increased invasiveness, possible infection dissemination, and increased morbidity and mortality, which poses a great challenge for the clinical treatment of *Klebsiella pneumoniae*. The detection rate of hv-CRKP is high in CRKP, and most of them are nosocomial infections. It is necessary to strengthen the monitoring and control of multi-drug-resistant bacteria and hypervirulent strains, further enhance the implementation of nosocomial infection prevention and control measures, prevent the spread of nosocomial infections, improve the clinical cure rate and safeguard medical health of patients.

## References

[1] Li W, Sun G, Yu Y, Li N, Chen M, Jin R, et al. Increasing occurrence of antimicrobial-resistant hypervirulent (hypermucoviscous) klebsiella pneumoniae isolates in China. Clin Infect Dis 2014;58:225–32. 10.1093/cid/cit675.

[2] Li J, Ren J, Wang W, Wang G, Gu G, Wu X, et al. Risk factors and clinical outcomes of hypervirulent klebsiella pneumoniae induced bloodstream infections. Eur J Clin Microbiol 2018;37:679–89. 10.1007/s10096-017-3160-z.

[3] Siu LK, Yeh K-M, Lin J-C, Fung C-P, Chang F-Y. Klebsiella pneumoniae liver abscess: A new invasive syndrome. Lancet Infect Dis 2012;12:881–7. 10.1016/S1473-3099(12)70205-0.

[4] Chen L, Kreiswirth BN. Convergence of carbapenem-resistance and hypervirulence in klebsiella pneumoniae. Lancet Infect Dis 2018;18:2–3. 10.1016/S1473-3099(17)30517-0.

[5] Wu H, Li D, Zhou H, Sun Y, Guo L, Shen D. Bacteremia and other body site infection caused by hypervirulent and classic klebsiella pneumoniae. Microb Pathog 2017;104:254–62. 10.1016/j.micpath.2017.01.049.

[6] Tian D, Liu X, Chen W, Zhou Y, Hu D, Wang W, et al. Prevalence of hypervirulent and carbapenem-resistant klebsiella pneumoniae under divergent evolutionary patterns. Emerg Microbes Infect 2022;11:1936–49. 10.1080/22221751.2022.2103454.

[7] Liu C, Dong N, Chan EWC, Chen S, Zhang R. Molecular epidemiology of carbapenem-resistant klebsiella pneumoniae in China, 2016–20. Lancet Infect Dis 2022;22:167–8. 10.1016/S1473-3099(22)00009-3.

[8] Munoz-Price LS, Poirel L, Bonomo RA, Schwaber MJ, Daikos GL, Cormican M, et al. Clinical epidemiology of the global expansion of klebsiella pneumoniae carbapenemases. Lancet Infect Dis 2013;13:785–96. 10.1016/S1473-3099(13)70190-7.

[9] Wei Z-Q, Du X-X, Yu Y-S, Shen P, Chen Y-G, Li L-J. Plasmid-mediated KPC-2 in a klebsiella pneumoniae isolate from China. Antimicrob Agents Chemother 2007;51:763–5. 10.1128/AAC.01053-06.

[10] Sader HS, Castanheira M, Shortridge D, Mendes RE, Flamm RK. Antimicrobial activity of ceftazidime-avibactam tested against multidrug-resistant enterobacteriaceae and pseudomonas aeruginosa isolates from U.S. medical centers, 2013 to 2016. Antimicrob Agents Chemother 2017;61:e01045–17. 10.1128/AAC.01045-17.

[11] Wangchinda W, Laohasakprasit K, Lerdlamyong K, Thamlikitkul V. Epidemiology of carbapenem-resistant enterobacterales infection and colonization in hospitalized patients at a university hospital in thailand. Infect Drug Resist 2022;Volume 15:2199– 210. 10.2147/IDR.S361013.

[12] Lin Y-T, Siu LK, Lin J-C, Chen T-L, Tseng C-P, Yeh K-M, et al. Seroepidemiology of klebsiella pneumoniae colonizing the intestinal tract of healthy Chinese and overseas chinese adults in asian countries. BMC Microbiol 2012;12:13. 10.1186/1471-2180-12-13.

[13] Choby JE, Howard-Anderson J, Weiss DS. Hypervirulent klebsiella pneumoniae - clinical and molecular perspectives. J Intern Med 2020;287:283–300. 10.1111/joim.13007.

[14] Parrott AM, Shi J, Aaron J, Green DA, Whittier S, Wu F. Detection of multiple hypervirulent klebsiella pneumoniae strains in a new york city hospital through screening of virulence genes. Clin Microbiol Infec 2021;27:583–9. 10.1016/j.cmi.2020.05.012.

[15] Zhao Q, Guo L, Wang L, Zhao Q, Shen D. Prevalence and characteristics of surgical site hypervirulent klebsiella pneumoniae isolates. J Clin Lab Anal 2020;34:e23364. 10.1002/jcla.23364.

[16] Yang Y, Liu J-H, Hu X-X, Zhang W, Nie T-Y, Yang X-Y, et al. Clinical and microbiological characteristics of hypervirulent klebsiella pneumoniae (hvKp) in a hospital from north China. J Infect Dev Ctries 2020;14:606–13. 10.3855/jidc.12288.

[17] Hwang J-H, Handigund M, Hwang J-H, Cho YG, Kim DS, Lee J. Clinical features and risk factors associated with 30-day mortality in patients with pneumonia caused by hypervirulent klebsiella pneumoniae (hvKP). Ann Lab Med 2020;40:481–7. 10.3343/alm.2020.40.6.481.

[18] Russo TA, Olson R, MacDonald U, Metzger D, Maltese LM, Drake EJ, et al. Aerobactin mediates virulence and accounts for increased siderophore production under iron-limiting conditions by hypervirulent (hypermucoviscous) klebsiella pneumoniae. Infect Immun 2014;82:2356–67. 10.1128/IAI.01667-13.

[19] Catalán-Nájera JC, Garza-Ramos U, Barrios-Camacho H. Hypervirulence and hypermucoviscosity: Two different but complementary klebsiella spp. phenotypes? Virulence 2017;8:1111–23. 10.1080/21505594.2017.1317412.

[20] Holt KE, Wertheim H, Zadoks RN, Baker S, Whitehouse CA, Dance D, et al. Genomic analysis of diversity, population structure, virulence, and antimicrobial resistance in klebsiella pneumoniae, an urgent threat to public health. Proc Natl Acad Sci 5 2015;112:E3574–3581. 10.1073/pnas.1501049112.

[21] Yeh K-M, Kurup A, Siu LK, Koh YL, Fung C-P, Lin J-C, et al. Capsular serotype K1 or K2, rather than magA and rmpA, is a major virulence determinant for klebsiella pneumoniae liver abscess in singapore and taiwan. J Clin Microbiol 2007;45:466–71. 10.1128/JCM.01150-06.

